# Caecal epithelium-derived thymic stromal lymphopoietin is not required for protective immunity against whipworm

**DOI:** 10.1101/2025.10.10.681633

**Authors:** Connor P. Lynch, Seona Thompson, Erin-Claire Pallott, Laura Campbell, Richard K. Grencis

## Abstract

Recent research examining classical type two alarmin TSLP has uncovered non-epithelial cellular sources of the cytokine during various murine challenges. *Trichuris muris* is a key model of type two immunity in the intestine with expulsion being TSLP-dependent. Previous research was suggestive of epithelial production of TSLP in the caecum, but it’s production during infection has not been characterised in detail. Here, using qPCR, transcript-based imaging, and conditional knockout, we demonstrate caecal epithelium to be a poor source of *Tslp*, and that non-epithelial upregulation occurs during *Trichuris* infection, peaking around day 14 post-infection. Conditional knockout of TSLP expression in caecal epithelial cells did not here impact parasite expulsion, but, surprisingly, neutralisation of TSLP from days 15 to 25 produced full susceptibility. These data build on previous work to raise important questions regarding the interaction of TSLP with immunity broadly, as well as the conditions for production of immunity to intestinal helminths.

## Introduction

Thymic stromal lymphopoietin (TSLP) is a four alpha helix bundle cytokine, emerging early in vertebrate evolution as a paralog of lymphocyte growth factor IL-7 [1], with orthologs in both humans and mice. A key cytokine in promoting the type two immune response against the intestinal helminth *Trichuris muris* (*Tm*), whipworm [2], TSLP has been a molecule of substantial interest in the field of human atopy since first observed to produce a model of murine asthma when overexpressed in the lung [3], with TSLP-neutralising tezepelumab having shown promising phase III clinical trial results in the treatment of severe asthma [4, 5]. Despite this, key aspects of TSLP’s biology remain unclear: despite being broadly characterised as an epithelial alarmin [6–10], recent work in mice has indicated murine *Tslp* expression to be fibroblast-restricted in the small intestine [11] and macrophage restricted in the skin [12], during the challenges examined (feeding and *Leishmania* infection, respectively). Additionally, both papers characterise the alarmin as acting directly on type two innate lymphoid cells to enable their function, in contrast to research suggesting TSLP action on dendritic cells [13, 14]. Although the level of conservation between murine and human TSLP functionality is not clear in light of the short inhibitory form of TSLP expressed by humans [15] but seemingly not by mice, human lung macrophage production of TSLP in response to stimulus *ex vivo* has been observed [16], suggestive that non-epithelial expression of TSLP in various murine tissues may shed light on unstudied TSLP sources in humans. Additionally, the effectiveness of tezepelumab in treating human asthma, in which adaptive immune responses to allergens would be overwhelmingly long established, would suggest TSLP as primarily functioning to enable lymphoid effector cell production of pro-asthmatic cytokines, rather than acting on APCs.

*Tm* is an orally-transmitted, caecum-dwelling gastrointestinal helminth and a closely related species to the widespread human-infecting *Trichuris trichiura*. While low level infection results in type one adaptive immune responses and chronic infection, high level infections (high doses of eggs, HD) result in type two adaptive immunity and IL-13 mediated epithelial expulsion of the parasite from the caecum [17, 18]. HD *Tm infection* is therefore a useful model of the necessary conditions for induction of type two immunity, and among helminths, TSLP appears uniquely necessary for supporting these type two immune responses in this system [14]. This makes HD *Tm* infection an ideal model for studying natural induction of TSLP, and examination of how TSLP contributes to a functional immune response leading to parasite expulsion.

Here, using conditional knockout models, we demonstrate that caecal epithelial cells are poor sources of *Tslp* both during homeostasis and HD *Tm* infection, and that caecal epithelial cell-conditional TSLP knockout does not impact immunity to HD *Tm* infection. We also observe that TSLP exerts its impact on immunity predominantly during the third and fourth weeks post-infection, consistent with publications from other groups suggesting a role for TSLP in licensing adaptor cell function in tissues.

## Results

### *Tslp* is poorly expressed by the caecal epithelial compartment and not substantially upregulated in the caecum during HD *Tm* infection

To examine any changes in expression of *Tslp* by epithelial cells during HD *Tm* infection, the epithelial compartment was isolated from the caecal lamina propria and submucosa of naïve male C57BL/6 mice, and alongside mesenteric adipose tissue associated with the caecum and the combined mesenteric lymph nodes, the resultant samples analysed for *Tslp* expression via RT-qPCR (**Fig. 1b**). Adipose tissue has been recently observed to produce *Tslp* during *H. polygyrus* bakeri infection [19], and here exhibited the highest naïve expression level of *Tslp* among tissues sampled. Flow cytometry analysis indicated expansion during HD *Tm* infection of adipose stromal cells resembling that observed by Kabat *et al*, however qPCR data from adipose and ELISA of isolated mesenchymal adipose tissue progenitor cells indicated no significant increase in *Tslp* expression or production at d14 p.i. (S.Fig 1). The epithelial samples contained on average approximately 8-fold lower levels of *Tslp* transcript than the lamina propria samples (also containing the submucosa and muscularis layers), 10-fold lower than lymph node samples, and 20-fold lower than adipose tissue. Next, *Tslp* expression in whole caecal samples at various timepoints during HD *Tm* infection, including a secondary infection administered at d39, a timepoint at which the primary infection is completely expelled (**Fig. 1c**). No significant changes in expression were measured during primary infection, but a trend towards decreased expression at days35 and and during secondary infection achieved significance at day 47, with a significant ∼3-fold decrease, perhaps attributable to changes to the caecum during infection such as increased caecal patch size and epithelial layer expansion, and suggestive of no tissue-wide upregulation of *Tslp* during infection. Next, to compare the magnitude of alarmin responses with another intestinal helminth infection, samples from the ileum following *Trichinella spiralis* (*Ts*) infection and the caecum following HD *Tm* infection were subjected to RT-qPCR for *Tslp* alongside *Il33* and *Il25* (**Fig. 1d-i**). All alarmins exhibited trends towards increased expression during *Ts* infection, ∼7-fold for *Tslp*, ∼5-fold for *Il33*, and ∼9-fold for *Il25* by day 10 p.i, when compared with naïve controls. However, no *Tslp* or *Il33* increases were observed in the HD *Tm*-infected caecum. Additionally, *Il25* transcript was undetectable in the caecum, consistent with poor tuft cell counts in the caecum compared with the small intestine as observed previously [20]. Overall, qPCR gave no indications of substantial tissue-wide changes to alarmin expression in the caecum during early HD *Tm* infection.

**Figure 1.**
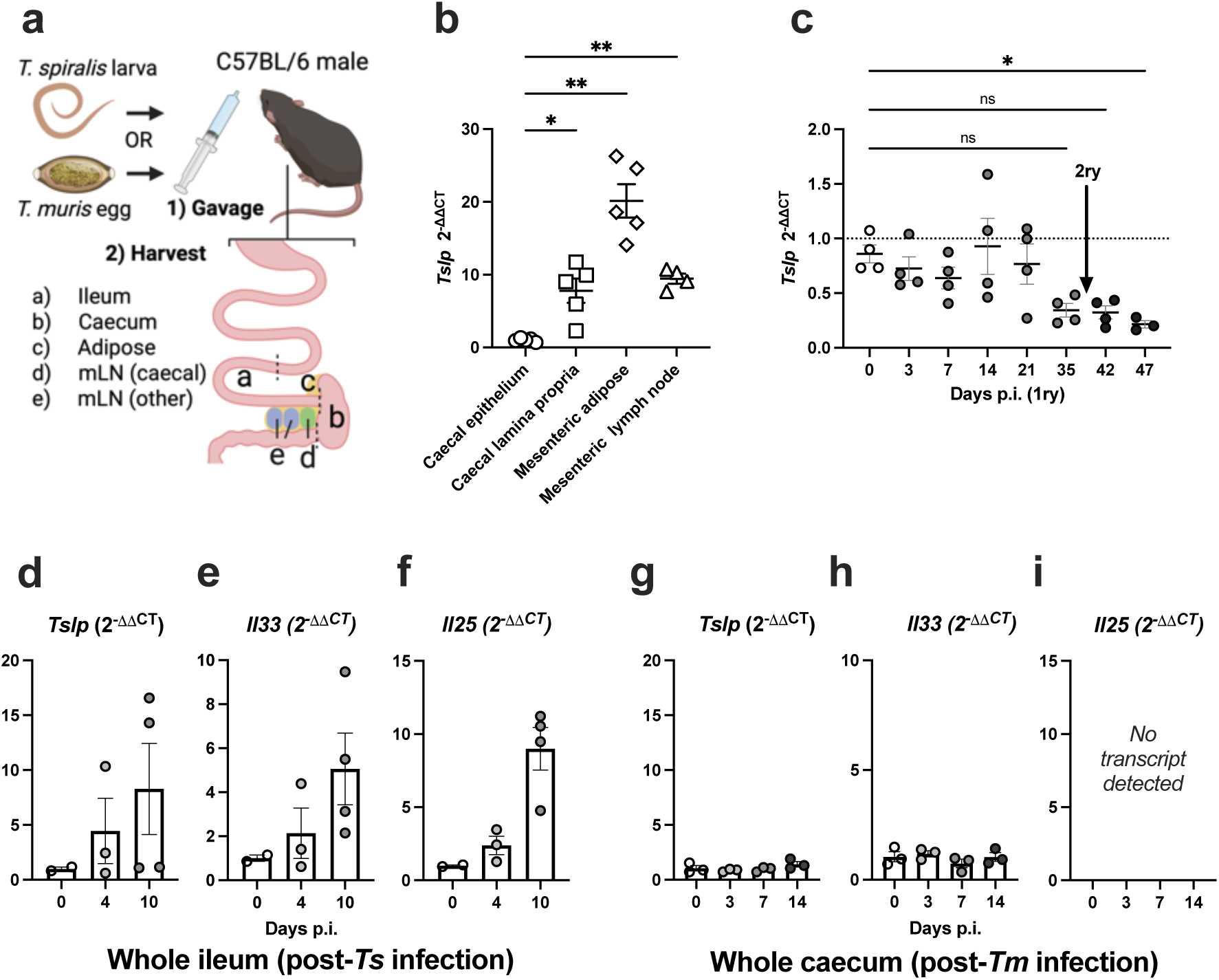
*Tslp* expression within caecal compartments and associated tissues, and tissue alarmin expression during *Trichinella* and *Trichuris* infection. (**a**) Overview of the two infection protocols, with an anatomical guide to the tissues harvested. (**b**) Epithelium was isolated from the caeca of naïve mice, RNA extraction performed, and cDNA reverse transcribed as described in. qPCR was performed targeting *Tslp* and *Rpl13* transcripts, and *Rpl13* CT subtracted from *Tslp* CT for each sample to generate ΔCT values shown (mixed-effects analysis testing, n=4-5 mice per group). (**c**) Mice infected with 300 *Tm* eggs on day 0 (and again on day 39), and their caeca harvested and processed for RT-qPCR using given timepoints, with ΔCT values again generated against *Rpl13*, ΔΔCT values generated against day 0ΔCT, and log transformed to give fold-change (2^-ΔΔCT^). (Kruskal-Wallace testing, n=3-4 mice per group). (**d-i**) Whole iliac intestinal tissue was harvested from *Ts*-infected mice at various timepoints p.i., was examined for alarmin transcription *via* qPCR, targeting *Tslp* (**d**), *Il33* (**e**), and *Il25* (**c**) transcripts (Kruskal-Wallace, n=2-4 mice per group). (**g-i**) Whole caecal tissue from HD *Tm* (300-egg dose) infected mice was similarly examined at different timepoints (Kruskal-Wallace, n=3 mice per group). All samples were generated from male C57BL/6 mice between 8 and 16 weeks old.

### Caecal epithelial knockout of TSLP does not replicate the susceptibility to high dose *Trichuris* infection produced by systemic TSLP neutralisation

In light of the surprising absence of either *Tslp* upregulation during HD Tm infection, and the poor expression by caecal epithelial cells, we opted to produce a conditional knockout mouse to remove expression of TSLP in caecal epithelial cells, and examine any impairment of the immune response to HD *Tm* infection. Observations of the Villin^β-gal^ mouse from our lab previously suggested relatively poor expression of Villin in the caecum (data not shown), commonly used in Cre-flox systems in epithelial cell knockouts in the small intestine [21]. To verify this, we examined a publicly available single cell sequencing dataset of the murine caecal epithelium for expression of *Vil1*, *Car1*, as well as *Epcam* [22]. In terms of both mean transcript count per epithelial cell as well as percentage of positive cells, *Car1* exhibited higher expression in caecal epithelium than *Vil1*, and was thus selected as the more suitable Cre promotor site (**Fig. 2a-b**). A Car1-Cre TSLP(flox/flox) mouse was created, in which exon 2 of the *Tslp* gene is flanked by LoxP sequences, and Cre recombinase enzyme is expressed under the Car1 promotor, constitutively expressed by proximal colonic and caecal epithelial cells, causing excision in *Car1*+ cells of genomic DNA encoding TSLP exon 2, rendering functional TSLP protein untranscribable in this cell type (**Fig. 2c**). Although Car1 is expressed by mast cells in addition to epithelial cells [23], mast cells appear to play little role in expulsion of HD *Tm* [24], limiting any caveats regarding off-target effects. TSLPΔCAR1 mice, as well as Cre-negative littermates (TSLPΔWT), were infected with 300- or 200-egg doses and adult parasites in the caecum counted at day 35 p.i. (**Fig. 2a**). 200-egg dose was included as a more sensitive model of TH2 cell induction, given the correlation between higher dose and stronger type two immunity. No differences were observed in expulsion between TSLPΔCAR1 and TSLPΔWT mice in either experiment, with complete expulsion of the 300-egg dose (**Fig. 2d**), and equivalent partial expulsion of the 200-egg dose between groups (**Fig. 2e**). To verify the previously reported requirement for TSLP signalling in expulsion of HD *Tm* infection [2, 14], a 300-egg infection was repeated with the addition of a group of littermates treated with anti-TSLP antibody every five days throughout infection, starting at d0. In this group, all mice remained infected by d35 p.i., while untreated TSLPΔCAR1 mice and littermates expelled all parasites (**Fig. 2f**). To validate Cre recombinase expression in the TSLPΔCAR1 mouse, epithelial cells of the caecum were isolated from the caeca of naïve TSLPΔCAR1 mice and littermates, and qPCR performed to measure Cre recombinase RNA expression: Cre recombinase RNA expression was observed in TSLPΔCAR1 epithelial samples to be >100-fold higher than in the lamina propria sample, or in either sample from TSLPΔWT mice (**Fig. 2g-h**). Expression level in TSLPΔCAR1 epithelial samples, measured by ΔCT versus *Rpl13* was in most samples comparable to that obtained from TSLPΔCAR1 genomic DNA from ear punches, (**Fig. 2i**) suggesting robust expression of the enzyme in the caecal epithelium in this knockout.

**Figure 2.**
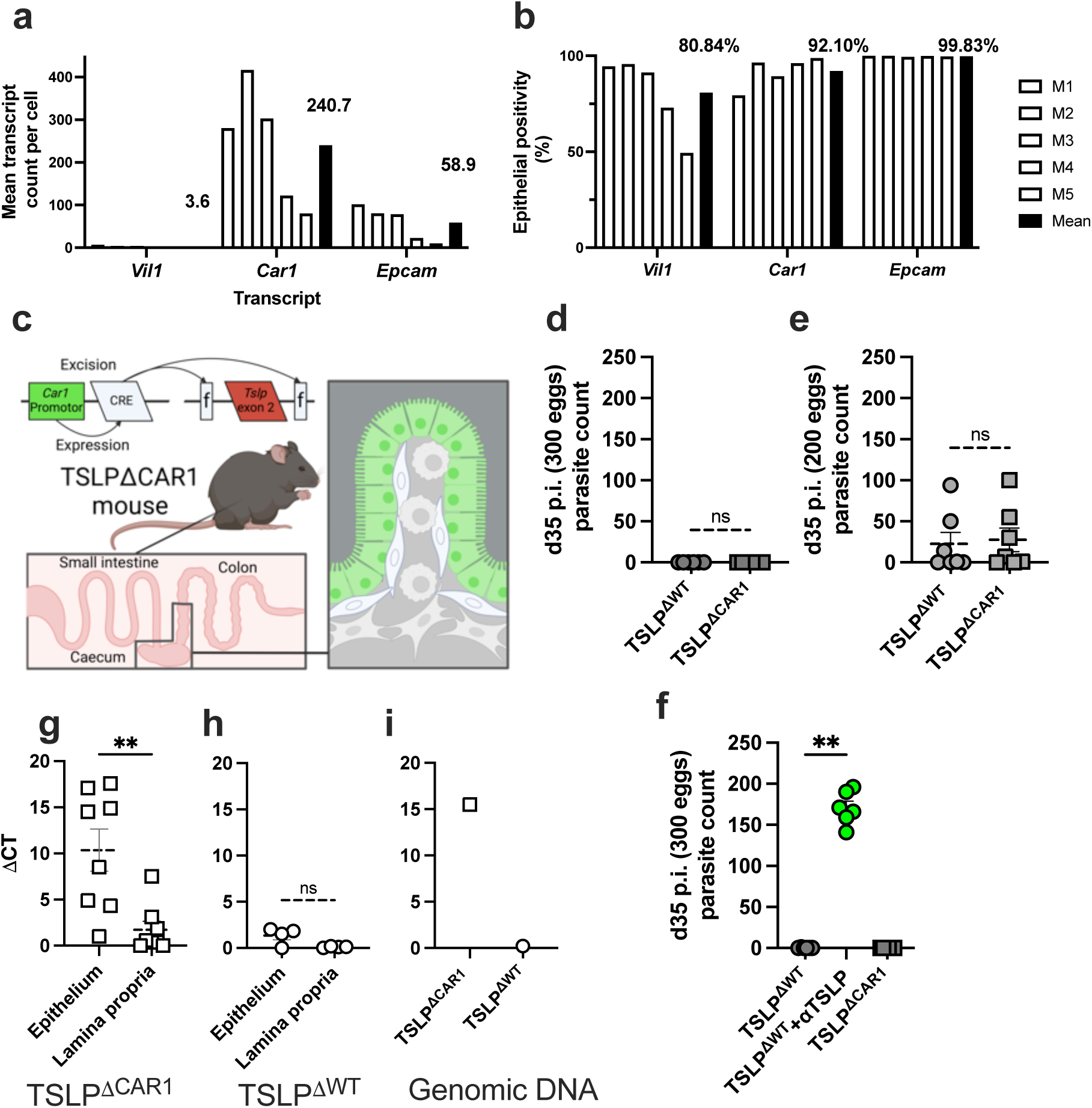
Validation and HD *Tm* susceptibility of the TSLPΔCAR1 caecal epithelial TSLP knockout mouse. (**a-b**) Single cell sequencing data from epithelial cells of caeca from five female C57BL/6 mice at 8 to 18 weeks old, produced as described here, with underlying matrix data deposited at GSE168448 in the NCBI GEO repository analysed via Microsoft Excel. Expression data graphed as percentage of positive cells (>0 transcripts) is given in **a**, while average transcript expression is given in **b** (Kruskal-Wallace testing). Mean bars are value labelled for each transcript. **c**) An overview of the TSLPΔCAR1 mouse knocking out *Tslp* expression in caecal and proximal colonic epithelial cells. Counts of adult parasites recovered from caeca in TSLPΔCAR1 and littermate control mice (TSLPΔWT) are shown, following 300-egg (**d**) and 200-egg (**e**) infections, at day 35 p.i. with *Tm* (Mann-Whitney testing, n=6-7 mice per group). (**f**) Groups and infection as in **d** repeated, plus a group of littermate control mice administered 100μg of anti-TSLP antibody *via* intraperitoneal injection every five days from d0 to d25 p.i. (Kruskal-Wallace testing, n=6 mice per group). *Cre recombinase* transcript ΔCT values generated against *Rpl13* housekeeping transcript, detected *via* RT-qPCR results from cDNA reverse transcribed from RNA isolated from enzymatically separated epithelial cells and remaining lamina propria (plus submucosa and muscularis layers), given for naïve TSLPΔCAR1 mice (**e**) and littermate controls (**h**) (Wilcoxon testing, n=7 mice per group). The same reaction was performed on genomic DNA from both groups in (**i**) (n=1 mouse per group). Mice between 12 and 16 weeks old at time of harvest.

### TSLP neutralisation, but not epithelial TSLP knockout of TSLP produces impaired IL-13-regulated epithelial responses to *Trichuris* infection

To gain insight into how immunity to HD *Tm* infection might be altered by impaired TSLP signalling, levels of parasite-specific antibody in serum and changes to the epithelial layer of the caecum were examined under several conditions: wild type littermates were separated into uninfected (TSLPΔWT naive) and day 35 post-infection (TSLPΔWT) groups, while epithelial TSLP knockouts were separated into groups treated with either isotype control antibody (TSLPΔCAR1+ISO, or “isotype treated”) or anti-TSLP antibody (TSLPΔCAR1+αTSLP, or “αTSLP treated”) every five days from days 0 to 25, prior to harvest at day 35 (**Fig. 3a**). Serum parasite-specific IgG1, used as a proxy readout of TH2 differentiation in the lymph node [25], was unimpacted by either epithelial TSLP knockout or TSLP neutralisation (**Fig. 3b**). TH1-associated IgG2a was likewise statistically not significant, despite a trend towards increase in the αTSLP treated group more in line with previously published data from TSLPR knockout mice [2] (**Fig. 3c**), suggesting only a marginal impact of TSLP neutralisation on B cell responses. Partial to total expulsion of parasites was observed in untreated littermates and isotype-treated epithelial knockouts, in contrast to the chronically infected anti-TSLP treated mice (**Fig. 3d**). Examining epithelial responses associated with IL-13-mediated parasite expulsion using H&E staining of tissue sections from the caecum, increased crypt length relative to naïve WT samples, indicative of chronic infection, was only observed in the αTSLP treated knockout mice (**Fig. 3e**), which likewise differed from infected wild type mice in having a reduced goblet cell count when normalised for changes to crypt length (**Fig. 3f-g**). Isotype-treated epithelial knockouts did not differ from either naïve or infected littermates in crypt length or indications of goblet cell hyperplasia, indicating no impairment of worm expulsion.

**Figure 3.**
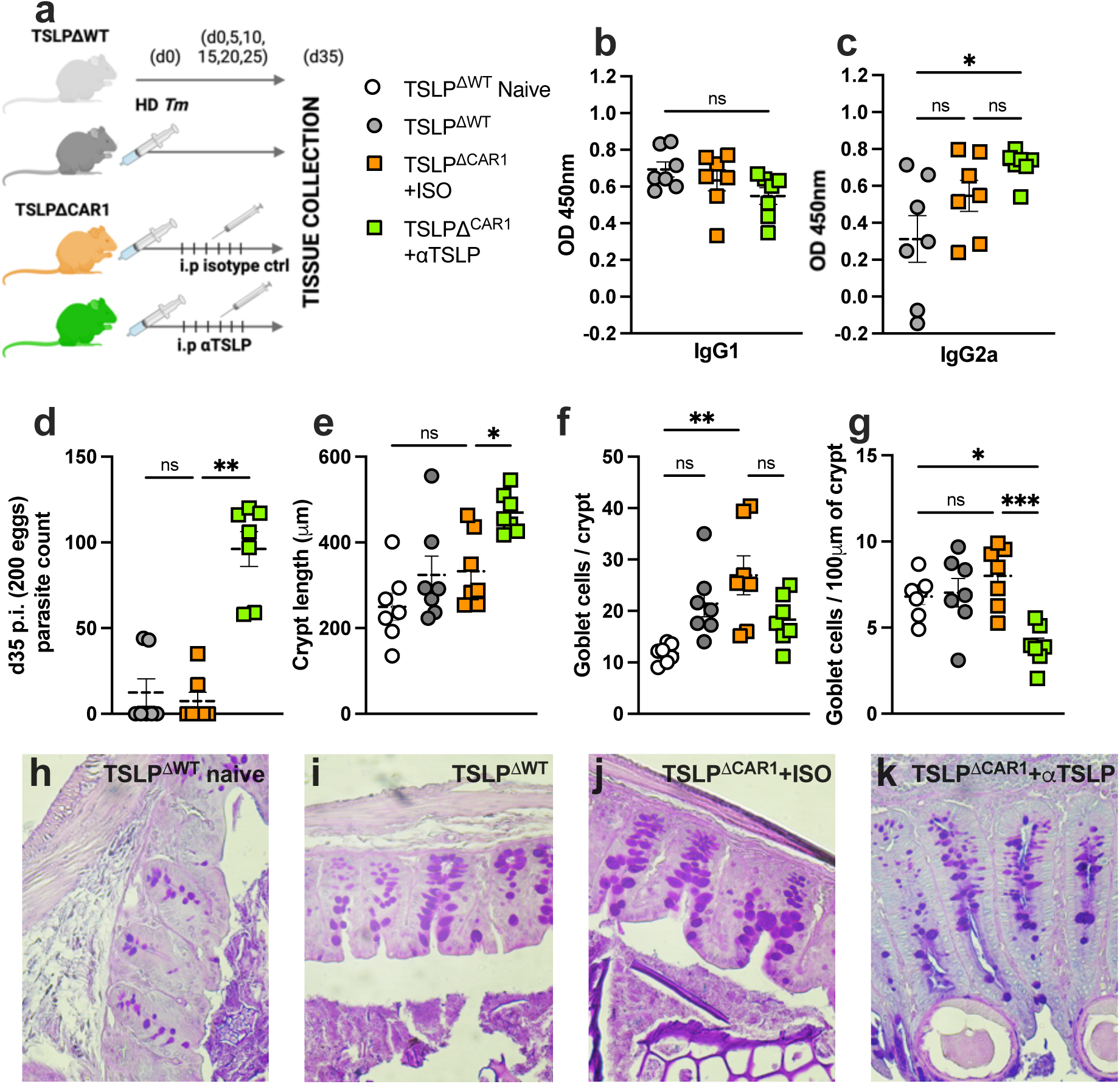
Immune readouts during *Trichuris* infection in epithelial conditional TSLP knockout and systemically TSLP neutralised mice. (**a**) Experiment overview: TSLPΔCAR1 mice and littermate controls were infected with 200 *Tm* eggs, TSLPΔCAR1 mice being separated into an isotype control treated (rat IgG2a) group and an anti-TSLP (rat IgG2a) treated group, each receiving 100ug of antibody administered intraperitoneally every five days from d0-25 p.i., prior to sample collection at d35 p.i. (**b**) Counts of adult parasites recovered from caeca in each group. Serum level of parasite-specific IgG1a (**c**) and IgG2a/c (**d**), measured *via* ELISA at 1 in 20 dilution and graphed in arbitrary optical density units, for infected groups (a-c Kruskal-Wallace testing). Length of caecal crypts (**e**), number of goblet cells per crypt (**f**), and goblet cells per 100ul of crypt (**g**) were quantified for all groups *via* analysis of H&E histological staining in ImageJ (Ordinary one-way ANOVA). (**h-k**) Representative images of H&E sections from each group used in quantification. Each point used in **d-f** represents an average of at least six crypts from one mouse. n=7 mice per group, between 12 and 16 weeks old at time of harvest.

### *In situ* hybridisation indicates non-epithelial upregulation of *Tslp* in the caecum during high dose *Trichuris* infection

RNAscope was used to visualise expression of Tslp mRNA in the caecum throughout HD *Tm* infection. Tslp+ cells were detected at all timepoints, including naïve mice (**Fig. 4a**), localised to rare lamina propria- and submucosa-resident cells (**Fig. 4b-e**). Expression in epithelial cells appeared to be negligible, with cells directly in contact with the intestinal lumen rarely exhibiting more than 1 transcript. This was corroborated by co-staining with EpCAM antibody, showing minimal expression of *Tslp* by EpCAM+ epithelial cells in both naïve and infected (d14 p.i.) mice (**Fig. 4g-i**). Quantification of both the number of *Tslp* transcripts and the number of *Tslp* positive cells (>3 transcripts per nucleus, thresholding determined from random probe control staining) indicated no significance increase in expression, due the rarity of positive crypts, but a trend towards maximum expression by both metrics in the lamina propria around day 7 (**Fig. 4i**) and in the submucosa at day 14 p.i (**Fig. 4j**).

**Figure 4.**
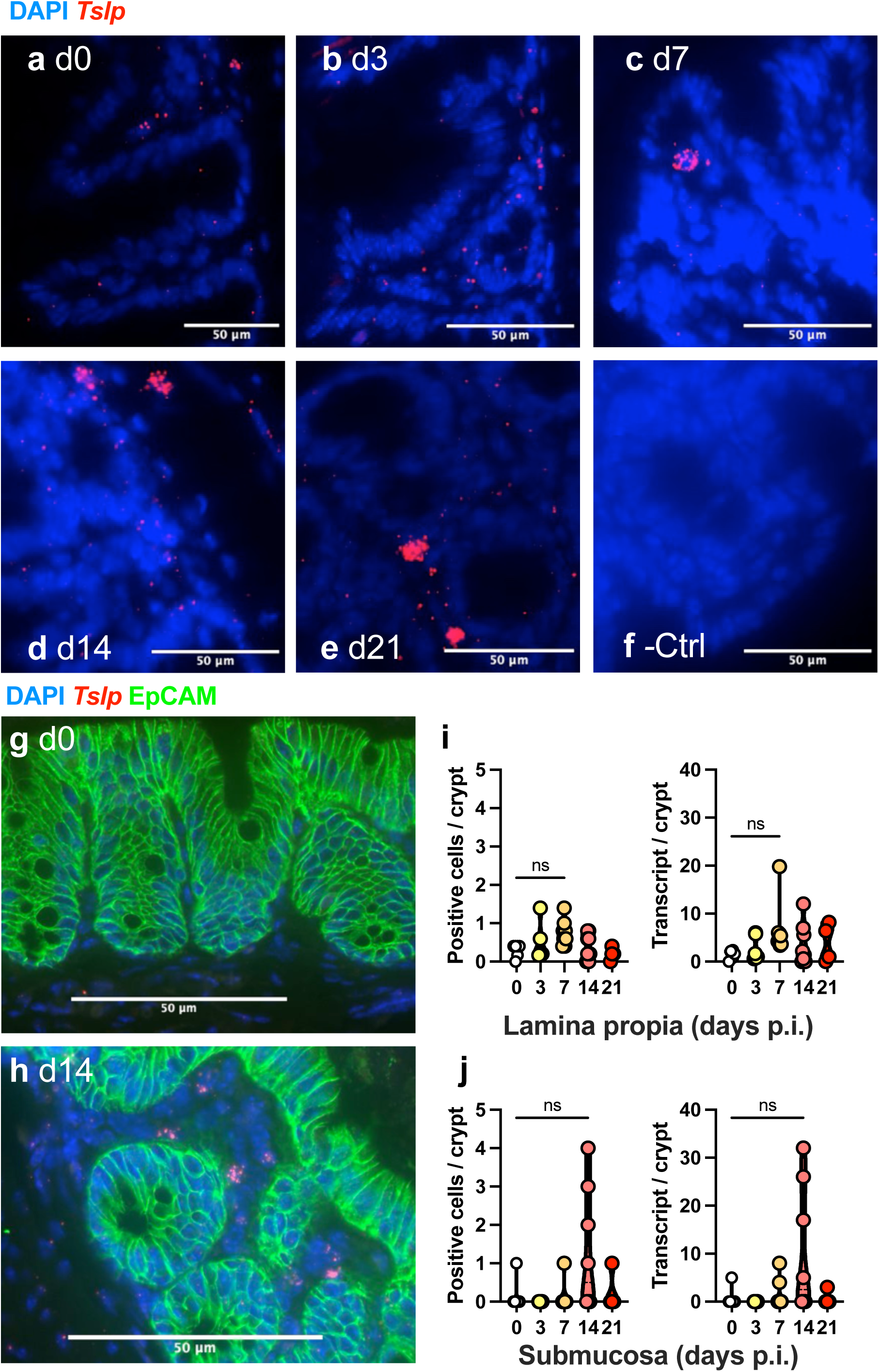
**(previous page) *Tslp* transcript imaging and quantification in the HD Tm-infected caecum**. Representative images of RNAscope-stained FFPE sections from male C57BL/6 mouse caeca post-HD *Tm* (300-eggs) infection in naïve mice (**a**), and mice at days 3 (**b**), 7 (**c**), 14 (**d**), and 21 (**e**) post-infection. *Tslp* transcript (in red) counterstained with DAPI for nuclei (blue) in sections represented by **a-e**, with a random oligomer probe substituted for *Tslp* in negative control sections (**f**). **g-h**) The same samples were used to generate sections stained similarly, with the addition of EpCAM antibody co-staining, with representative images from the naïve (**g**) and day 14 post-infection (**h**) sections displayed. Quantification performed on RNAscope-stained sections for *Tslp* transcript count per crypt as well as positive cells per crypt, for the lamina propria (**c**) and submucosa (**d**) (Kruskal-Wallace, n=4-6 mice per group). Positive cells were defined as ≥3 transcripts overlaid on a nucleus. Each point used in d-f represents an average of at least four crypts from one mouse, from three separate infections. Mice were between 12 and 16 weeks old at time of harvest.

### TSLP contributes to *Trichuris* expulsion primarily from day 14-post infection onwards

Considering the surprisingly late upregulation of Tslp during infection, a timed neutralisation experiment was performed in which one group of mice received anti-TSLP treatment at days 0, 5, and 10 p.i. (early), while another group was treated at days 15, 20, and 20 (late), alongside an untreated group (**Fig. 5**). Surprisingly, while the early treated group exhibited parasite burdens at d35 p.i., late treated mice were significantly more heavily infected (>2-fold higher parasite burden) (**Fig. 5b**). This was accompanied by an increased crypt length (**Fig. 5e**) and decreased goblet cells per micron of crypt (**Fig. 5c-d**) in late treated mice versus untreated controls (**Fig. 5e**), indicative of chronic infection and impaired IL-13-mediated goblet cell hyperplasia, respectively. In these epithelial readouts, the early treated group exhibited non-significant changes intermediate between untreated and late treated mice, potentially suggestive of administered antibody persisting into late infection. Unusually, late treated mice also showed increased parasite-specific IgG1 in serum (**Fig. 5f)**, normally associated with TH2 immunity and parasite expulsion. IgG2a levels, associated with chronic infection, were largely unimpacted in both groups (**Fig, 5g**).

**Figure 5.**
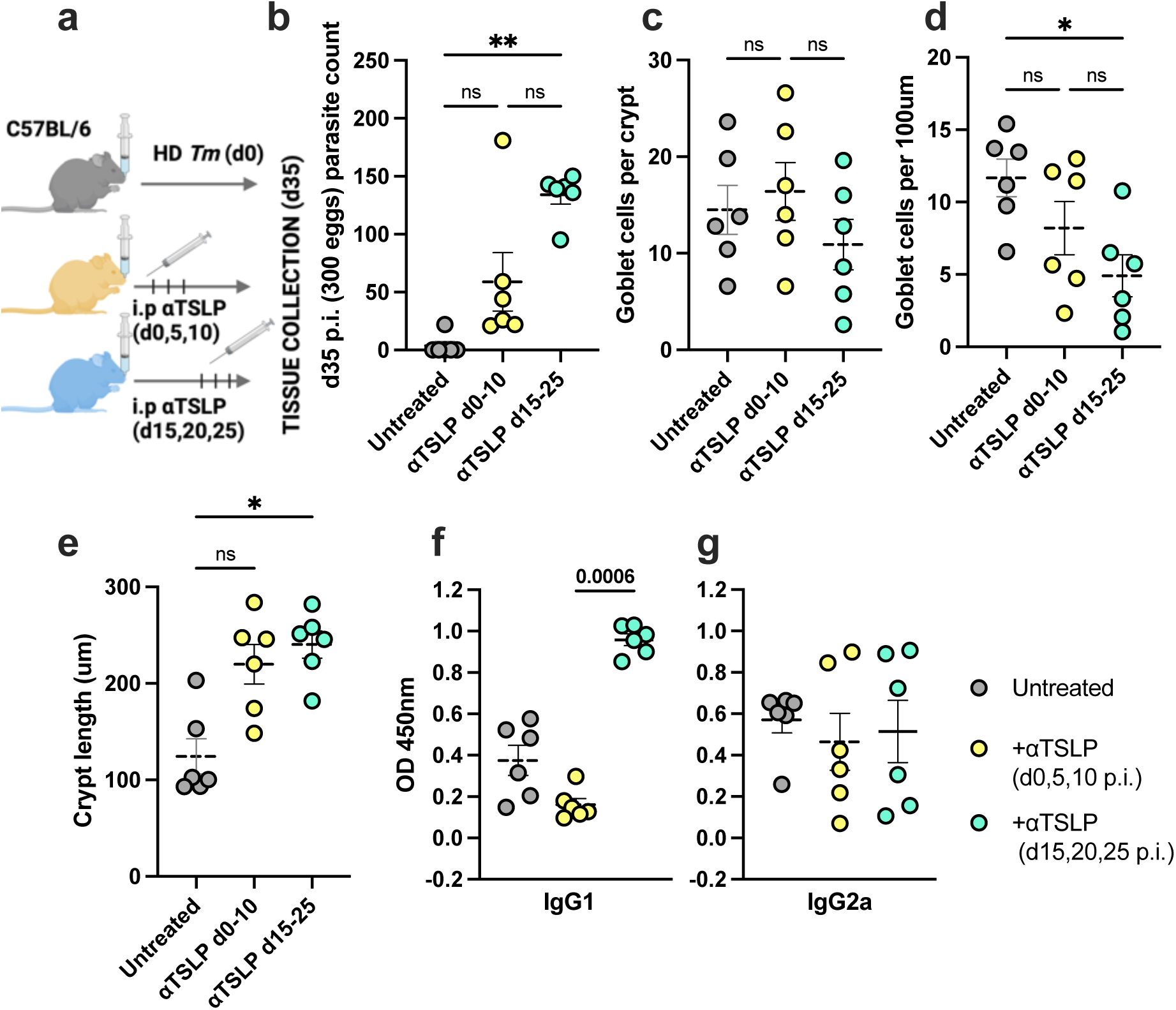
Impact of early and late TSLP neutralisation on parasite expulsion, antibody responses, and stromal immunity following HD *Tm* infection. (a-b) C57BL/6 mice and were infected with 400 *Tm* eggs and treated with 100ug of anti-TSLP antibody intraperitoneally in two different dosage regimes, as well as an untreated group, harvested at day 35 p.i. **a**) Experimental summary. **b**) Counts of adult parasites recovered from caeca in each group (Kruskal-Wallace). Number of goblet cells per crypt (**b**), goblet cells per 100ul of crypt (**d**), and length of crypts (**e**) were quantified for all groups *via* analysis of Alcian blue histological staining in ImageJ (Ordinary one-way ANOVA). Serum level of parasite-specific IgG1a (**f**) and IgG2a/c (**g**), was measured *via* ELISA and graphed in arbitrary optical density units (Kruskal-Wallace). Each point in (**c-e**) used in **f**-**h** represents an average of at least six crypts from one mouse. N=6 mice per group, between 12 and 16 weeks old at time of harvest.

## Discussion

Epithelial cells are frequently the first cellular host point of contact with pathogens and parasites, and for this reason are studied closely in the context of barrier immunity. Considering the IEC niche inhabited by *Tm* and a role TSLP in HD *Tm* expulsion, taken with the assumption that IECs are TSLP sources at other barrier sites, TSLP production by caecal IECs in response to contact with *Tm* has remained the null hypothesised role in immunity. However, to date this has proved difficult to further evidence mechanistically. Results indicating either successful antibody staining of murine TSLP, or demonstrating qPCR use to detect changes *Tslp* transcription, are both uncommon. One study observed a 4-fold increase in whole caeca from BALB/c mice at day 7 p.i. with HD *Tm* [26], another study isolating epithelial cells observed a trend towards higher TSLP production *via* qPCR at both 24hrs and 7 days p.i. [27]: whether this occurs in C57BL/6 mice has not been demonstrated, and may represent a significant difference between the strains which contributes to their differing susceptibilities to *Tm* infection . Other research has, however, observed TSLP expression in the caecal epithelial layer in C57BL/6 mice *via* antibody staining [2], and a subsequent study posited dendritic cell detection of epithelium-produced TSLP as the key TSLP contribution to HD *Tm* immunity [13], although epithelial TSLP expression *in vivo* was not directly observed here. A recent single cell sequencing examination of the caecum in early infection with HD *Tm* (days 1 and 3 p.i.) uncovered no changes in *Tslp* expression, and minimal expression in IECs [28]. Overall, *in vivo* evidence for caecal IEC *Tslp* upregulation during HD Tm infection, or functional contribution to immunity, is minimal. Data presented here indicates *Tslp* transcript to be poorly expressed in the epithelial compartment relative to other caecal compartments, and, surprisingly, no substantial increase in transcription was observed in the caecum during infection. Normal melt curves and prior publications using these primers to detect alarmin changes during *Hp* infection [29] both indicated primer functionality, and although results were unexpected, previous studies observing *Tslp* expression in the caecum did not isolate the epithelial compartment [2] as performed here. This could be due to *Tslp* transcription being highly localised, upregulated only in crypts contacting parasites, diluting any observable increases during analysis of whole tissue: this would be consistent with the observable, but not statistically significant, increases in *Tslp* expression observed via *in situ* hybridisation. The significant decrease observed in *Tslp* expression during secondary infection may be due to expansion of the caecal patch, a lymphoid structure at the caecal tip, reducing the mass of normal caecal parenchyma sampled.

Interestingly, various recent papers have recently documented TSLPR expression on ILC2s and TH2 cells [11, 12, 19, 30–34] to a far greater extent than has so far been demonstrated for dendritic cell detection of TSLP *in vivo*. This is consistent with a theorised model of alarmin function in which TSLP acts to license effector cell cytokine production in tissues, rather than polarising adaptive immune cell generation via antigen presenting cells [35], which is also consistent with data here indicating that TSLP contributes to immunity to HD *Tm* infection during the expulsion phase. The decrease in goblet cell numbers per crypt below even naïve levels in infected and anti-TSLP treated mice suggests that homeostatic TSLP may be supporting homeostatic IL-13 production which promotes goblet cell differentiation.

Homeostatic *Tslp* production by Foxl1+ cells, observed here *via* RNAscope, would be concordant with recent work suggesting feeding-stimulated *Tslp* production by these cells promoting ILC2 IL-13 production at homeostasis. Data here, indicating expansion of Tslp expression during infection in Foxl1-cells, would suggest a separate Tslp induction mechanism is at play during HD *Tm*, While antibody responses were, as might be expected, essentially identical between TSLPΔCAR1 and littermates, the impact of anti-TSLP treatment on parasite-specific antibody responses was surprisingly minor. Unusual antibody responses were also observed during the phased neutralisation experiments, in which highly susceptible mice administered anti-TSLP during late infection were found to produce high levels of parasite-specific IgG1. This may be the result of a normally resistant strain of mouse retaining a high parasite burden in spite of normal lymph node responses in the late-treated group, resulting in a heightened response due to increased parasite antigen load or intestinal damage. Although intact antibody responses is consistent with a role for TSLP in licensing lymphoid cell effector functions in tissue, rather than polarising APCs, recent interesting work has noted a role for TSLP in enabling T-B cell crosstalk [36]. Other recent work examining follicular helper T cell responses suggests a broadly pro-TH1 signature in these cells following *Tm* infection, in contrast to *Hp* infection [37], in consensus with previous work indicating a limited role for B cell-produced antibodies in directly shaping protective immunity to HD *Tm* infection [24, 38]. As a result, the effects of anti-TSLP treatment of antibody responses may be distinct from the susceptibility to infection it produces.

Recently published data [11] using a TSLP reporter mouse model is consistent with our observations of non-epithelial cells being TSLP producers following HD *Tm* infection: their observations indicated a sub-epithelial fibroblast subset, *Foxl1+* telocytes, as key TSLP producers in the intestine following feeding. Published data from our lab observed likely co-expression of *Foxl1* and *Tslp* in the caecum, but indicated that increases in caecal *Tslp* expressing cells in the lamina propria and submucosa do not appear to occur in *Foxl1*+ cells. *Ccl24*+ macrophages, demonstrated as key TSLP producers in the skin during *Leishmaniasis* [12], were similarly observed to produce Tslp in the caecum, and expand Ccl24 transcription during infection, but did not produce notably higher levels of *Tslp* during infection. Moreover, conditional TSLP knockout from FOXL1+ or CCL24+ cells did not change the resistant phenotype following HD Tm infection, contraindicating either cell type as the critical *Tslp* upregulating cells observed here via RNAscope. In the absence of an effective immunofluorescence-compatible antibody targeting TSLP, spatial transcriptomic and proteomic approaches which permit imaging of TSLP+cells would aid identification and characterisation of the critical cells, and a deeper study of their potential interaction with neighbouring cells e.g. lymphoid effector cells *in vivo*. While such a mechanism is only hypothetical, it is strongly supported by recent research indicating TSLP-mediated type two lymphoid cell activation *in vivo* [11, 12, 19, 30, 33, 39] . Here, the concept of TSLP responses in the intestine as being epithelial, and as contributing to immunity early on in infection prior to the expulsion phase, is not supported, and creates space in the field for characterisation of a new mechanism by which TSLP influences the immune system, with important implications for diseases at mucosal sites in which TSLP plays an important role, such as allergic asthma. The level of conservation between murine and human TSLP functionality is not clear, considering the short inhibitory form of TSLP expressed by humans [15] but seemingly not by mice. Recent single cell sequencing data from the human lung observed expression of *Il33* by epithelial cells, but not *Tslp* [40]. Another study of human lung samples observed a correlation between epithelium-expression EGR1, Il33 expression, and asthma, while no correlation between EGR1 and *Tslp* observed in epithelial cells. Human lung macrophage production of TSLP in response to stimulus *ex vivo* has been observed [16], however, suggestive that recently observed non-epithelial expression of TSLP in various murine tissues may shed light on unstudied TSLP sources in humans. Significant gaps remain in the field regarding the specific mechanism by which TSLP, among other alarmins contribute to the immunology of asthma: detailed investigation along the lines of inquiry followed here would be highly informative in this area of study.

## Acknowledgements

We would like to thank the Manchester Genome editing unit and the Biological Services Facility at the University of Manchester for their contributions to this work. Thanks to Dovydas Širvinskas *et al* for producing the open access data used in Figure 2. Anti-TSLP neutralising antibody was generously donated by Amgen Inc.

## Methods

### Mice

C57BL/6J.*Tslp*Em1Uman mice (TSLP-floxed, or TSLP^f/f^) mice were generated by the University of Manchester Genome Editing Unit, the two loxP sites being integrated in the intergenic region upstream (5’) of the gene and within the *Tslp* intron between exons 2 and 3 (3’), using a CRISPR-Cas9 system to integrate long single stranded DNA, generated as described by [41], administered *via* pronuclear microinjection of cryopreserved 1-cell embryos alongside an sgRNA (Sigma-Aldrich, g295 – gaggctctccccgcttagag, g330 – ttttacatgccaaatgtgtg), which were implanted into surrogate psuedopregnant mice after overnight culture of the embryos.

Genotyping was performed with REDExtract-N-Amp Tissue PCR Kit (Sigma-Aldrich) to validate insertions *via* gDNA ear punches in weaned mice. C57BL/6 mice were purchased from Charles River or bred in house at the University of Manchester Biological Services Unit (derived initially from Envigo C57BL/6J). All mice were maintained in a specific pathogen free (SPF) facility at the University of Manchester, in a 12:12 hr light:dark cycle at 21±5°C and 55±10% humidity. Mice were housed in cages of one to six, with rodent chow and sterile water provided *ad libidum*. All mice were euthanised *via* rising concentration of CO_2_. Experiments were performed in accordance with the United Kingdom Animals (Scientific Procedures) Act of 1986, under project licenses P043A082 (2021-2023) and PP0172300 (2024-2025) and were subject to local ethical review by the University of Manchester Animal Welfare and Ethical Review Body (AWERB) and followed ARRIVE 2.0 guidelines. The mice utilized in these experiments were not randomized but cages were randomly assigned to different treatment groups for experiments. All mice used for experiments were between 8 and 16 weeks of age. Knockout mouse experiments made use of both sexes of mice, while C57BL/6 experiments used male mice only. Details of the ages and sexes of mice used for each experiment are provided in figure legends.

#### Trichuris muris

The Edinburgh strain of *T. muris* was used in all experiments originally obtained from The Wellcome Research Laboratories, London, and is routinely passaged within our laboratory. For infections, mice were administered between 200 and 400 eggs *via* oral gavage in Milli-Q water, using eggs between 60 and 75% infectivity, as measured *via* batch test infection of SCID immunodeficient mice, bred in house at the University of Manchester (strain originally obtained from the Fox Chase Cancer Centre, Philadelphia). For parasite count data, infected mice were euthanised and caeca and proximal colon harvested. Tissue was opened longitudinally and contents washed out with water, prior to counting of adult parasites by eye using the MZ75 dissection microscope (Leica). For E/S acquisition, X-linked severe combined immunodeficient (SCID) mice were infected with 200 *T. muris* eggs by oral gavage, and their caeca harvested at day 42 p.i. into RPMI-1640 (Sigma-Aldritch) plus 500 U/mL penicillin and 500ug/mL streptomycin (Sigma Aldritch) (RPMI+P/S). Mice were euthanised, and caeca stored in pre-warmed (approx. 40°C) RPMI were extracted from the caecum using fine forceps and incubated in RPMI+P/S for 4hr at 37°C, with the media being harvested and replaced for a second 16hr incubation. Both lots of media were centrifuged in 50mL tubes (Falcon) to pellet eggs, supernatant was collected into a new 50mL tube for E/S production.

Pelleted eggs were resuspended in Milli-Q (Sigma-Aldrich) water, passed through a 70μm filter to remove adult parasites and intestinal debris, and embryonated *via* incubation in T75 culture flasks (Corning) for 6-8 weeks. Embryonated eggs were transferred to 4°C storage for use in infecting mice, after infectivity testing to ensure >50% infectivity (i.e. 200 egg administration to a mouse producing >100 embedded larvae at a timepoint prior to expulsion, d12-16 p.i.). The supernatant separated from eggs for E/S production was first filtered through a 0.22μm syringe filter (Sartorius). Concentration was performed using Amicon Ultra centrifuge units, the resulting media centrifuged at 3000xg for 15min at 4°C. Resulting media was dialysed against PBS (pH 7.4, Gibco) for 24 hr at 4°C, then the protein concentration of the E/S was determined using a Nanodrop (NanoDrop One, Thermo Fisher). Resulting E/S was aliquoted and stored at −20 °C until use.

### *In vivo* administration of antibodies

Treated mice were injected intra-peritoneally with 100μl of 1mg/mL rat-anti-mouse anti-TSLP (Amgen, IgG2a not commercially available) every five days on alternating sides. Isotype control-treated mice received 100μl of isotype control antibody (BioXcell, cat#BE0085) at the same timepoints.

### Anti-parasite immunoglobulin ELISA

Flat-bottomed 96-well plates (Corning) were coated with 5μg/mL of parasite E/S diluted in sodium carbonate bicarbonate buffer (pH 9.6), and incubated overnight at 4°C. The following day, plates were washed five times (all wash steps were performed with 200ul per well of PBS plus 0.05% Tween-20 (Sigma), using the AquaMax 4000 (Molecular Devices).), and incubated with 100ul of 3% BSA (Sigma) in PBS for 1hr at room temperature (RT). Block was tipped from plates, and 100ul of serum dilution (top row 1:20, diluted 1:3, serially) was added to each well, and incubated for 90min at RT. Plates were washed three times, and incubated with biotin-conjugated αIgG1 (cat#MCA336B, BioRad) or αIgG2a (cat#553388, BD Biosciences), at 1:1000 in PBS for 1hr at RT. Plates were washed, and incubated with 1:1000 streptavidin-POD conjugate (Roche) for 1hr at RT. Plates were washed, and incubated with a 1:1 solution of development substrates A and B from the BD OptEIA TMB substrate reagent set (BD Biosciences). Wells were allowed to develop before a stop solution was added (50ul of 2M sulphuric acid).

Plates were read at 405nm using a VersaMax plate reader, and after verification of normal dilution:OD relationships for samples, a concentration of 1:120 was graphed for each sample on the plates.

### Histological staining and quantification

Caecal tip tissue dissected from mice were fixed by immersion in 4% neutral buffered formalin for 16hr, followed by transfer into 70% ethanol (Fisher) and wax embedding. Sections were cut at 5μm thickness *via* microtome (Leica RM2255) and mounted onto Superfrost Plus slides (Epredia).

Hemotoxylin and eosin (H&E) staining was performed as follows: 2min in running water, 4mins in Harris’ hematoxylin (Electron Microscopy Sciences), 10 seconds in 1% HCl in ethanol (acid-alcohol), 3mins in dH_2_O, 30sec in tap water, 2mins in dH_2_O, 30sec in 1% eosin, then rehydrated *via* ethanol gradient and mounted with Pertex and cover slips. Periodic acid-Schiff (PAS)/Alcian blue staining was performed as follows: 5min in 1% Alcian blue solution + 3% acetic acid (Sigma) if using Alcian blue protocol, 1min in dH_2_O, 5min in 1% periodic acid (Thermo Fisher, 1min in dH_2_O, 5min in tap water, 1min in dH_2_O, 15min in Schiff’s reagent (Sigma) if using PAS protocol, 1min in dH_2_O, 5min in tap water, 1min in dH_2_O, 30sec in Meyer’s haematoxylin (Sigma), 5min in tap water, then rehydrated *via* ethanol gradient and mounted with Pertex and cover slips. Imaging was performed using an Axioimager M2 upright microscope (Zeiss). Images were processed and analysed using Fiji/ImageJ. Goblet cell counts per crypt and crypt length data were generated in Fiji by for each mouse by counting the number of goblet cells (stained purple in H&E staining or blue in Alcian blue or PAS staining) per crypt crypt, measuring the crypts length, using 6-20 crypts per mouse to generate each graphed data point.

### RNAScope staining

RNAscope staining (RNAscope Multiplex Fluorescent Reagent Kit v2, Biotechne) was performed on tissues fixed and sections as described above, according to manufacturer’s instructions. Briefly, following digestion with protease and hydrogen peroxide, and treatment in antigen retrieval buffer, sections were stained using the Co-Detection Ancillary Kit with an anti-EpCAM antibody (cat# ab71916, Abcam) for 2hr at RT. Sections were washed and then hybridised with *Tslp* or negative control probes (targeting a nonsense sequence) detection probes at 40°C. Probe sequences are proprietary. Sections were treated with amplification reagent, fluorophores conjugated (Opal™ 570, Akoya Biosciences) stained with a goat anti-rabbit AlexaFluor 488 secondary antibody (cat#A-11008, Invitrogen) for 90min at RT, counterstained with DAPI, and mounted for image analysis. Imaging was performed using an Axioimager M2 upright microscope (Zeiss). Images were processed and analysed using Fiji/ImageJ.

### RNA extraction and cDNA synthesis

Tissue harvested from mice directly, or processed as described in **Enzymatic digestion of caecum**, prior to being added to 1mL of TRIzol (Thermo Fisher) in a 2mL lysing matrix D beat bead mill-compatible tube (MP Biomedicals). TRIzol stored tissues were thawed and homogenised using the Bead Mill 24 (Fisher Scientific) at 20Hz for 4min, then placed on ice. TRIzol stored cell suspensions were homogenised *via* pipetting prior to storage at -80°C. RNA extraction was performed using the Direct-zol RNA Microprep Kit (Zymo Research). Briefly, TRIzol was added to a column and centrifuged at 14,000xg for 30sec, washed with provided buffer, the RNA precipitated into the column matrix and repeatedly washed with buffer, before being eluted into 0.5mL tubes (Eppendorf), washed with 75% ethanol, air dried after centrifugation, and solublised in nuclease-free water (Invitrogen). Sample RNA concentration was determined using the NanoDrop One (Thermo Fisher), and concentration adjusted downwards with nuclease-free water if necessary. RNA was then stored at -80°C until reverse transcription could be performed. Reverse transcription reactions were performed using the Luna RT Supermix Kit (NEB), with the mass of RNA reverse transcribed kept consistent between samples within experiments. The resulting cDNA was stored at -20°C.

### qPCR

qPCR reactions were performed on cDNA samples, produced as earlier described, using LunaScript Universal qPCR Master Mix (NEB) in 96-well V-bottomed 0.2mL qPCR plates (ABI Fast Systems, Starlab). In short, 1μL of sample was combined with 0.5μL of forward primer, 0.5μL of reverse primer (both at 10mM), 4ul of LunaScript master mix, and 14μL of nuclease-free water. Samples were run in triplicate, and the mean value taken for analysis, using three or two replicates (divergent single replicates were excluded if >1 CT difference from replicates was measured, or abnormal melt curves or amplification plots noted). Amplification was performed using the QuantStudio3 thermocycler (Thermo Fisher). Analysis was performed using the Thermo Fisher Connect Platform to visualise melt and amplification curves, Microsoft Excel for calculation of ΔCT and ΔΔCT values. ΔCT values were calculated using *Rpl13* transcript as a housekeeping gene, tested in lab as producing more consistent CT values than alternatives *Bact* or *Gapdh* (data not shown). Melt curves and amplification plots were examined for all samples to validate primer binding and normal amplification. Calculation of 2^-ΔΔCT^ values used for graphing and analysis was performed by calculation of ΔΔCT values by subtraction of the control group’s mean CT from each individual experimental group CT, and calculating 2 to the power of this value’s negative, to produce a “fold change” in transcript levels between control and experimental samples.

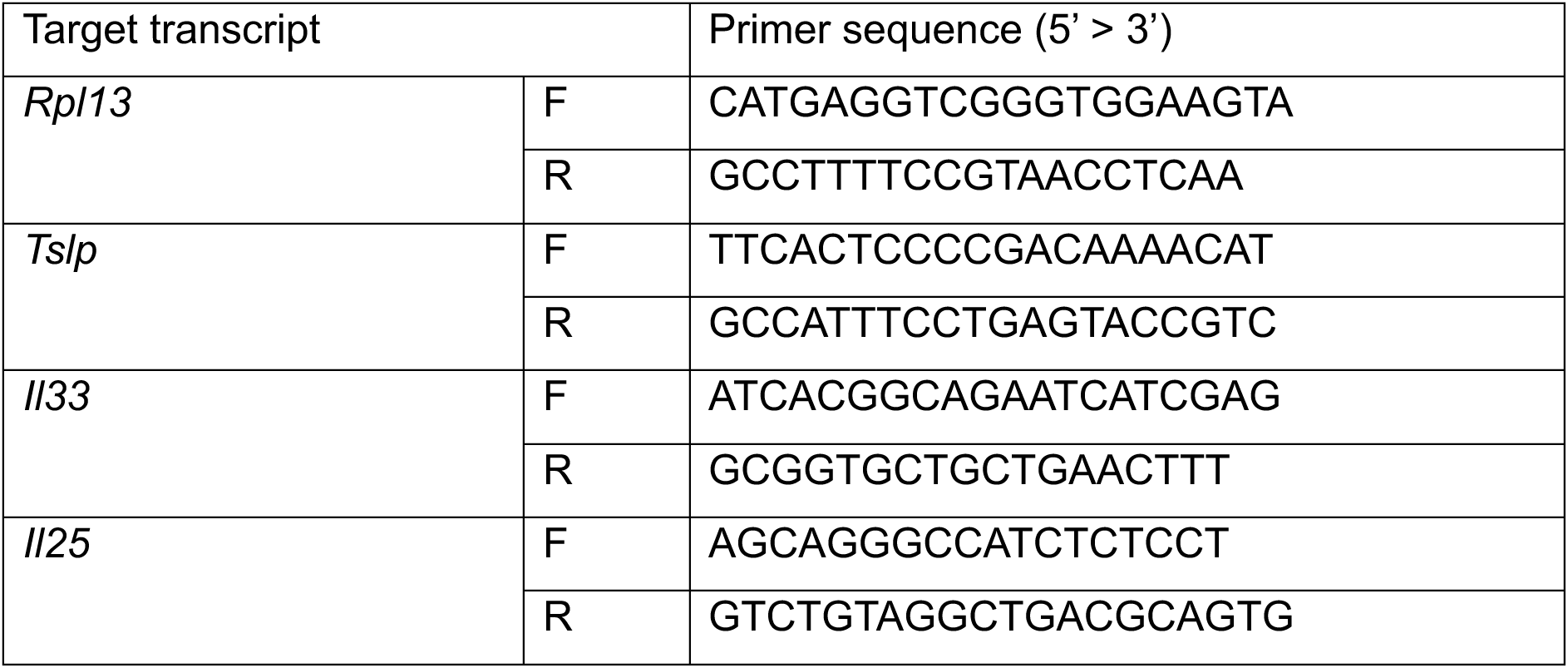

### Enzymatic digestion of caecum

The caecum was dissected from mice, the contents scraped out, and the tissue washed with chilled HBSS -Ca/Mg (without Ca/Mg) (Sigma) by flushing followed by shaking in a 10mL Falcon tube. Tissues were incubated in HBSS (-Ca/Mg) +5mM EDTA (Sigma) +1mM DTT (Sigma) for 30 minutes at 37°C in a shaking incubator, the buffer replaced halfway through the incubation. Tissues were then vortexed for 30sec then passed through a 100μm filter to isolate lamina propria from epithelial compartment single cell suspensions. At this point, the epithelial compartment and lamina propria were either transferred to separate lysing matrix D tubes for RNA extraction (see **RNA extraction and cDNA synthesis),** or the epithelial compartment discarded and lamina propria retrieved from the filter for further digestion. In the latter case, the lamina propria was minced with scissors and transferred into 5mL of digest buffer consisting of HBSS with calcium and magnesium (Gibco) + 1% FCS +1mg/mL of Liberase TL (Roche) +10μg/mL DNAse (Sigma Aldrich). Tissue was incubated for 45min at 37°C in a shaking incubator (mixed *via* pipetting at 15 and 30 min), then centrifuged at 400xg for 10min at 4°C. Cells were resuspended in cold PBS, filtered using a 70μm filter into a 10mL tube, and centrifuged again. Cells were resuspended in chilled PBS, transferred into a V-bottomed 96-well plate (Corning), and kept at 4°C on ice. Cells were then stained using the PrimeFlow assay.

### Enzymatic digestion of adipose tissue for flow cytometry

The adipose tissue attached to the caecum was dissected and stored in chilled fat media (low glucose DMEM (Sigma) +5g BSA (Melford) +25mL HEPES (1M) (Sigma). Samples transferred into digest buffer (per mouse, 3.5mL fat media + 0.7mg Liberase TL +0.875mg DNase at 37°C for 30min. 1.5mL of fat media + 5mM EDTA was added, tissue broken down by pipetting for 1min, then transferred through a 70µm filter into a 15mL tube and centrifuged at 400xg for 10min. Supernatant was removed, and cells resuspended in cold PBS for staining. Cells were stained in chilled PBS using Zombie UV™ Fixable *Via*bility Kit (BioLegend) at 1:1000 for 10min at 4°C in the dark. Cells were washed (400xg, 5min, 4°C) twice in chilled FACS buffer (PBS + 1% FCS), then stained using anti-mouse antibodies for 20 minutes at 4°C at 1:200 concentration in FACS buffer:

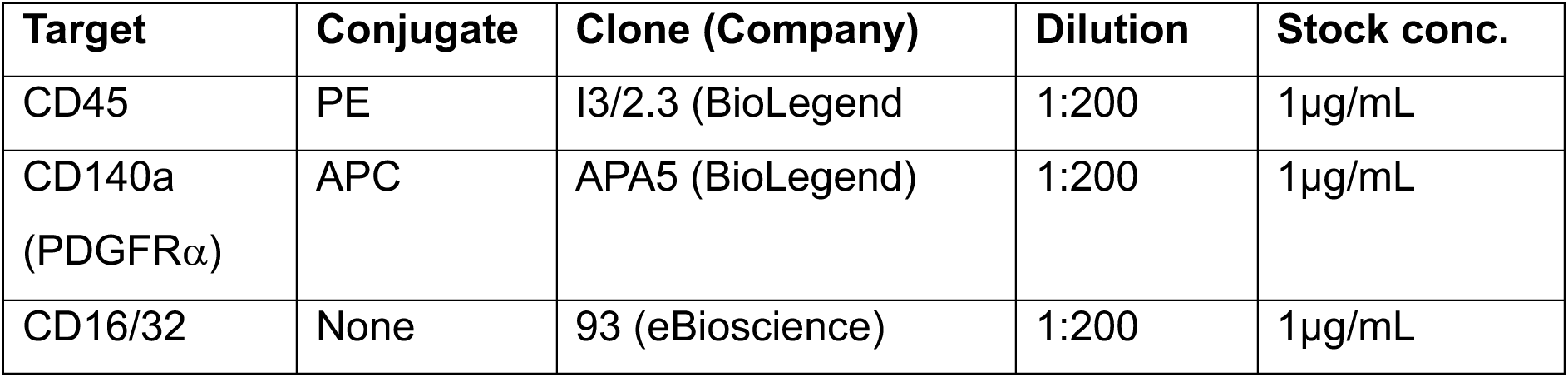

| Target | Conjugate | Clone (Company) | Dilution | Stock conc. |
| --- | --- | --- | --- | --- |
| CD45 | PE | I3/2.3 (BioLegend) | 1:200 | 1µg/mL |
| CD140a<br>(PDGFRα) | APC | APA5 (BioLegend) | 1:200 | 1µg/mL |
| CD16/32 | None | 93 (eBioscience) | 1:200 | 1µg/mL |

### Adipose tissue digest for MACS sorting, *ex vivo* culture, and ELISA

MACS sorting of adipose tissue was performed using the Adipose Tissue Progenitor Isolation Kit (Miltenyi Biotec) as per manufacturer’s instructions, using MidiMACS and MiniMACS separators. Sorted cells were plated at 10x10^5^ cells/mL for 24hr incubation at 37°C. Cells were transferred into 1.5mL tubes, centrifuged at 14,000xg for 10min at RT, and supernatant transferred into fresh 1.5mL tubes for ELISA analysis. ELISAs were performed using the Biotechne Duoset TSLP ELISA (cat#DY555-05), according to the manufacturer’s instructions, using supernatant from stimulated cell lines or *ex vivo* cells. OD values were acquired using the VersaMax plate reader.

