## Supplementary figures and images for "Caecal epithelium-derived thymic stromal lymphopoietin is not required for protective immunity against whipworm"

### Supplementary Figure 1

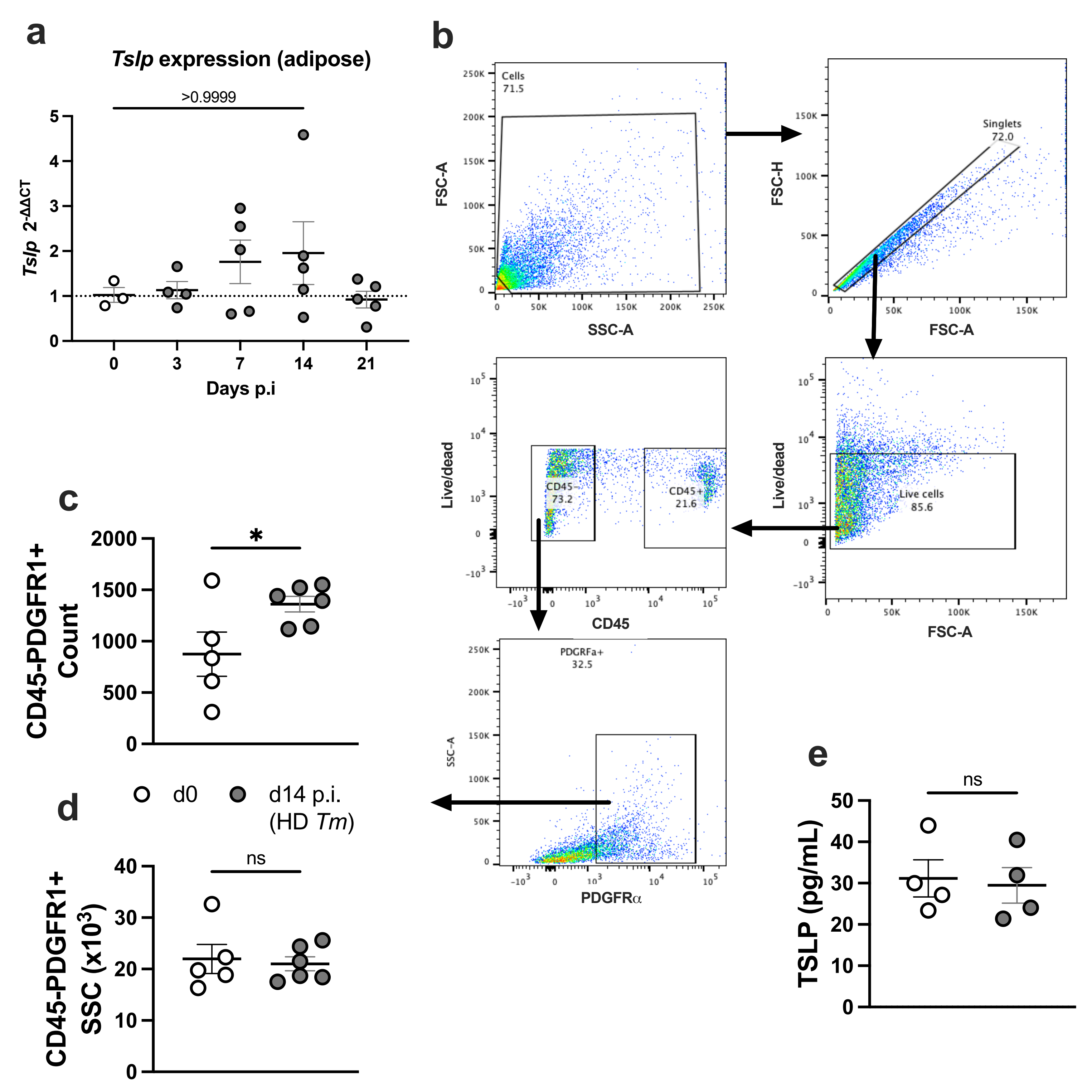
